# Longitudinal *in vivo* changes in radial peripapillary capillaries and optic nerve head structure in non-human primates with early experimental glaucoma

**DOI:** 10.1101/2021.07.11.450534

**Authors:** Gwen Musial, Suman Adhikari, Hanieh Mirhajianmoghadam, Hope M. Queener, Alexander W. Schill, Nimesh B. Patel, Jason Porter

## Abstract

**Purpose:** There is conflicting evidence as to whether a loss of radial peripapillary capillaries (RPCs) precedes neuronal loss in glaucoma. We examined the time-course of *in vivo* changes in RPCs, optic nerve head (ONH) structure, and retinal nerve fiber layer thickness (RNFLT) in experimental glaucoma (EG).

**Methods:** Spectral domain optical coherence tomography images were acquired before and approximately every 2 weeks after inducing unilateral EG in 9 rhesus monkeys to quantify mean anterior lamina cribrosa surface depth (ALCSD), minimum rim width (MRW), and RNFLT. Perfused RPC density was measured from adaptive optics scanning laser ophthalmoscope images acquired on the temporal half of the ONH. The time of first significant change was quantified as when values fell and remained outside of the 95% confidence interval established from control eyes.

**Results:** Mean ALCSD and/or MRW were the first parameters to change in 8 EG eyes. RPC density changed first in the 9th. At their first points of change, mean ALCSD posteriorly deformed by 100.2 ± 101.2 μm, MRW thinned by 82.3 ± 65.9 μm, RNFLT decreased by 25 ± 14 μm, and RPC density decreased by 4.5 ± 2.1%. RPC density decreased before RNFL thinning in 5 EG eyes. RNFLT decreased before RPC density decreased in 2 EG eyes, while 2 EG eyes had simultaneous changes.

**Conclusions:** In most EG eyes, RPC density decreased before (or simultaneous with) a change in RNFLT, suggesting that vascular factors may play a role in axonal loss in some eyes in early glaucoma.

## INTRODUCTION

Glaucoma is a group of eye diseases that are characterized by axonal loss and progressive retinal ganglion cell death, and can ultimately result in irreversible vision loss.^1^ Vascular-related factors have been shown to be associated with glaucoma, including lower diastolic perfusion pressure,^2–4^ migraines,^5,6^ nailfold capillary abnormalities,^7^ smoking,^4,8^ hypotension,^9,10^ and sleep apnea.^11,12^ Although vasculature has been considered an independent risk factor, vascular susceptibility to alterations in the translaminar pressure gradient are proposed to result in perfusion instabilities in capillaries within the lamina cribrosa and inner retinal tissues surrounding the optic nerve head (ONH).^13^ This imbalance gives rise to alterations in ocular blood flow and a reduced blood supply to ONH, laminar, and radial peripallary capillaries,^14–16^ potentially contributing to the pathogenesis of glaucoma.

The structural properties of the radial peripapillary capillaries (RPCs) may make them uniquely susceptible to damage resulting from elevated IOP in glaucoma. RPCs form the most superficial capillary bed in the retina and are potentially more fragile than other retinal capillaries, as they lack smooth muscle actin ensheathment, run in long parallel networks in the retinal nerve fiber layer (RNFL), and anastomose less frequently than other retinal capillaries.^14,15,17^ Consequently, RPCs possess less collateral blood supply and have fewer connections to adapt to necessary changes in autoregulation in glaucoma.

There is conflicting evidence as to whether RPCs play a role in retinal ganglion cell axon degeneration in glaucoma. Early work by Daicker et al.^18^ in donor eyes with chronic glaucoma and different optic nerve atrophies found no correlation between the distribution of atrophic RPCs and associated visual field defects. Subsequent histological work showed RPCs to be lost at the same rate as RNFL axons in experimental glaucoma^19^ and to not be significantly altered in their numbers in donor glaucomatous human eyes relative to normal eyes.^20^ However, the former of these two studies primarily evaluated the capillary volume within the ONH and not the RNFL. Conversely, other studies have reported significant losses of RPCs in experimental glaucoma^21^ and in donor eyes of glaucoma patients.^22^ More recent reports examining the RPC network *in vivo* using optical coherence tomography angiography (OCTA) found decreased peripapillary vessel density in glaucomatous eyes compared to age-matched controls^23–26^ and decreased peripapillary vessel density in the eyes of primary open-angle glaucoma patients with visual field defects.^27^ More recent work by Moghimi et al.^28^ found a weak tendency for eyes with lower baseline values of vessel perfusion density to have faster rates of RNFL thickness loss in mild to moderate glaucoma over a two-year period. In addition, studies have reported correlations between vessel density loss and visual field loss in the superotemporal and inferotemporal sectors around the ONH and in visual field maps.^25,26,29^ However, the general lack of *in vivo* data characterizing longitudinal changes in RPCs and retinal ganglion cell axons from a healthy state to early glaucoma has limited understanding of the relative time-course for when changes occur, as well as how differences in RPC, RNFL, and ONH geometries relate to disease progression in living eyes.

The primary purpose of this study was to determine whether changes in RPC perfusion occur prior to a loss of circumpapillary RNFL thickness in early experimental glaucoma. Longitudinal changes in RPC perfusion, ONH structure, and RNFL thickness were assessed using split detector adaptive optics^30,31^ and spectral domain optical coherence tomography imaging in living eyes of non-human primates with pressure-induced experimental glaucoma. Through the use of sensitive *in vivo* imaging techniques, this study better describes the sequence of early changes in capillary perfusion, ONH structure, and circumpapillary RNFL thickness in experimental glaucoma.

## METHODS

All animal care experimental procedures were approved by the University of Houston’s Institutional Animal Care and Use Committee and adhered to ARVO’s Statement for the Use of Animals in Ophthalmic and Vision Research. Nine adult rhesus macaques (mean age = 6.6 ± 1.2 years) were used in this study. Prior to inducing experimental glaucoma (EG), animals were imaged (as described below) to obtain baseline measurements. The trabecular meshwork of the right eye was scarred using a clinical 532nm laser (Zeiss Visulas 532; Carl Zeiss Meditec, Jena, Germany) to elevate the intraocular pressure of each monkey’s right eye (i.e., the experimental glaucoma eye), while the fellow eye served as a control.^32^ During these laser sessions, monkeys were anesthetized with ketamine (20–30 mg/kg) and xylazine (0.8–0.9 mg/kg). Multiple laser sessions, minimally separated by 2 weeks, were used to slowly build up and create sustained elevated pressure. The first session involved lasering 180° of the trabecular meshwork, and each subsequent session was limited to 90°.

Following the first laser session, animals were imaged approximately every 2 weeks throughout the duration of the experiment. Monkeys were anesthetized with 20–25 mg/kg ketamine and 0.8–0.9 mg/kg xylazine, and treated with a subcutaneous injection of atropine sulfate (0.04 mg/kg).^33^ Each monkey’s pupils were dilated with 2.5% phenylephrine and 1% tropicamide. A pharmacological agent (IOPIDINE; Alcon Laboratories, Inc., Fort Worth, TX, USA or COMBIGAN; Allergan, Inc., Irvine, CA, USA) was used at the start of each imaging experiment to reduce the animal’s IOP and best ensure that the values of parameters measured during the experiment were the result of chronic changes due to sustained elevation in IOP. IOP was measured using an applanation tonometer (Tono-Pen XL Applanation Tonometer; Reichert, Inc., NY, USA).

### BIOMETRIC SCALING

Biometric measurements of axial length, anterior chamber depth, lens thickness, and anterior corneal curvature were acquired from right and left eyes of each animal (LenStar; Haag-Streit, Köniz, Switzerland). These biometric parameters were used to convert field sizes in adaptive optics scanning laser ophthalmoscope (AOSLO) images from visual angle (in degrees) to physical retinal size (in micrometers). Conversions were performed by incorporating the measured biometry data into a 4-surface model eye.^34,35^

### SPECTRAL DOMAIN OPTICAL COHERENCE TOMOGRAPHY IMAGING

Scanning laser ophthalmoscope (SLO) fundus images and images of the ONH (15° or 30° field sizes) were acquired for each subject using spectral domain optical coherence tomography (SDOCT) (Heidelberg Spectralis HRA+OCT; Heidelberg Engineering, Heidelberg, Germany). These images assisted in navigating throughout the retina and optic nerve head (ONH) during AOSLO imaging sessions, as the AOSLO imaging field was small (approximately 2°). Monkey eyelids were held open using a lid speculum. Imaging was performed while monkeys wore a rigid gas permeable contact lens, which was used to prevent corneal dehydration and correct for any inherent spherical refractive errors.^34^ Control eyes were imaged prior to imaging the EG eye in each animal.

At baseline, SDOCT scans centered on the ONH were acquired to ensure good ocular health, and all subsequent scans were acquired using the instrument’s follow-up mode. Cross sectional radial scans (20°, 48 b-scans, high-resolution, 20 frame averaging) centered on the ONH were acquired with Enhanced Depth Imaging in all eyes. The inner limiting membrane (ILM) was automatically segmented in each b-scan using the SDOCT instrument’s software and any inaccuracies were manually corrected. Radial scans were exported from the SDOCT instrument and analyzed using programs written in MATLAB (MATLAB; The MathWorks, Inc., Natick, MA). The points of the retinal pigment epithelium (RPE)/Bruch’s membrane (BM) interface and anterior lamina cribrosa surface (ALCS) were manually marked in as many b-scans as possible.^35^ These delineated landmarks were used to calculate three ONH parameters. Bruch’s Membrane Opening (BMO) was calculated as the area enclosed by an ellipse best-fit to the marked BMO points in all radial scans. Mean anterior lamina cribrosa surface depth (ALCSD) was computed as the mean distance between a plane best-fit to the marked BMO points and a thin-plate spline surface that was fit to the marked ALCS points.^36^ Finally, mean Minimum Rim Width (MRW) was computed as the mean of the minimum distances between the marked BMO points and the ILM surface across all b-scans.^37,38^

RNFL thickness was measured from 12° circular scans centered on the ONH. The raw data was analyzed using a program written in MATLAB to determine the average RNFL thickness across the entire scan and average thickness in 60° sectors (as later described). The thickness of the RNFL was calculated as the distance between the automatically segmented ILM (following manual correction) and the manually marked, posterior boundary of the RNFL.^39^

### AOSLO IMAGING

High-resolution, *in vivo* measurements of RPC perfusion have been carried out in healthy and glaucomatous eyes primarily using optical coherence tomography angiography (OCTA) and AOSLO imaging. While OCTA has emerged as a powerful tool to image perfused vasculature (such as RPCs) at different retinal depths over a wide field,^24,40^ its relatively low lateral resolution (∼15 μm), due to the presence of ocular aberrations, limits its ability to image the smallest perfused retinal capillaries. Therefore, we used an AOSLO to correct the eye’s aberrations and provide high-resolution (∼2.5 μm) *in vivo* images of the finest capillary structures in the retina.

AOSLO images of perfused RPCs were acquired at baseline and in subsequent imaging sessions following the initial laser session. Each monkey’s head was positioned using a head mount attached to a 3-dimensional translation stage and was steered using the tip, tilt, and rotation capabilities of the head mount until the monkey’s ONH was within the AOSLO’s field of view. Through-focus AOSLO images were taken to determine the plane of best focus of the most superficial retinal vascular network (i.e., the RPCs in healthy eyes). *En face* reflectance videos of RNFL axon bundles were acquired using a confocal AOSLO imaging channel over a 2° field of view at a rate of 25 Hz using a superluminescent diode (SLD) light source (S-Series Broadlighter; Superlum, Carrigtwohill, Ireland) with a center wavelength of 840 nm (FWHM = 50 nm). The power of the SLD at the corneal plane was 150 µW, a value that was more than 10 times below the maximum permissible exposure for an imaging duration of 1 hour.^41^ Non-confocal, split detector adaptive optics videos^42^ were collected simultaneously with confocal videos at the same retinal location and depth, and were subsequently used to generate perfusion images.^43^ The 30° SLO image acquired with the Heidelberg Spectralis HRA+OCT served as a guide for navigating to the region of retina that was imaged with the AOSLO system. In each session, images of the RPC network were acquired in the temporal hemi-field around the ONH using the AOSLO system and were manually stitched together to form a montage of perfused vasculature. Images were first acquired in each animal’s EG eye before proceeding to image the contralateral control eye.

SLO images from all time-points were aligned in ImageJ (developed by Wayne Rasband, National Institutes of Health, Bethesda, MD; available at http://rsb.info.nih.gov/ij/index.html) for each subject using the plugin, StackReg, and were subsequently scaled based on each eye’s ocular biometry. Small modifications in image scale were made, if necessary, to match the scale of the RPC perfusion montage acquired with the AOSLO at the baseline time-point. Then, all subsequent RPC AOSLO montages were aligned to their corresponding SLO image taken at the same time-point. RPC montages were then segmented with a convolutional neural network (CNN) that has been previously described.^44^

### SECTOR ANALYSES

A 30° SLO image centered on the fovea was manually aligned with a 30° SLO image centered on the ONH (Adobe Photoshop; Adobe Systems, San Jose, CA). After manually marking the fovea, the fovea to Bruch’s Membrane Opening (FoBMO) axis^45^ was generated using a custom program written in MATLAB by connecting the manually marked foveal center with the center of the ellipse that was best-fit to the already marked BMO points (Fig. 1). The FoBMO axis served as the reference for generating 60° sectors around the ONH. In addition, circles were constructed at radial distances of 3° and 8° from the center of the BMO ellipse to form a 5° annulus over which RPC perfusion and RNFL parameters were analyzed.

**Figure 1.**
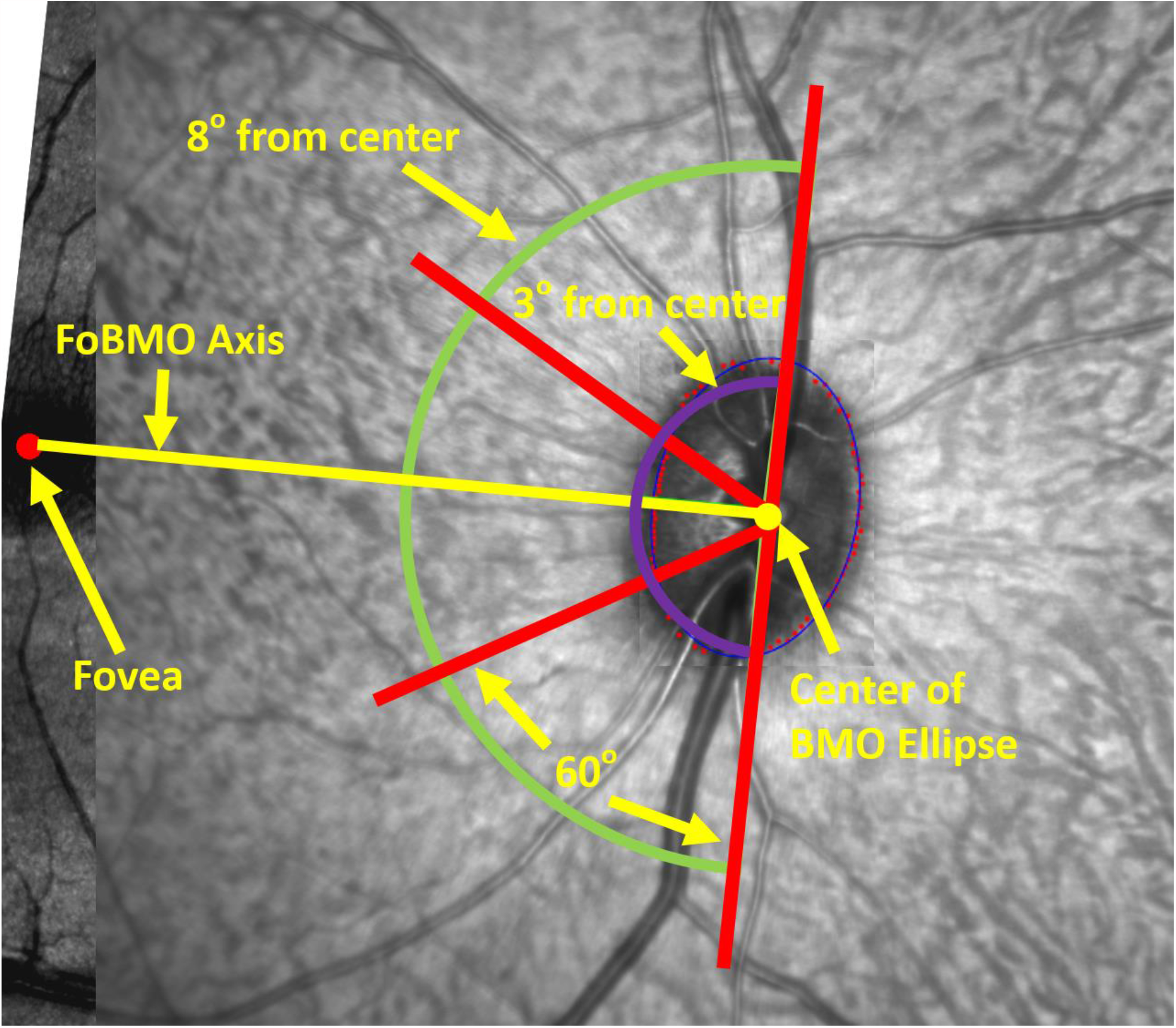
Use of the Fovea to Bruch’s Membrane Opening (FoBMO) axis for creating 60° sectors around the optic nerve head. An ellipse (thin blue line) was best fit to the manually marked BMO points (small red dots) and used to determine the center of the BMO (yellow dot). The foveal center was manually marked on the SLO image (large red dot) and connected to the center of BMO ellipse to form the FoBMO axis (yellow line). 60°sectors (corresponding to the angular subtense between red radial lines) were used in combination with annuli extended radially from 3° from the center of the BMO ellipse (innermost purple arc) to 8° from the center of the BMO ellipse (outermost green arc) for RPC analysis regions.

RPC density was calculated from the binary CNN segmentations as the percentage of pixels that contained perfused capillaries relative to the total area that was imaged and within a defined FoBMO sector (Fig. 1). For example, when analyzing the superior sector for a given session, a mask was made that was the union of the superior FoBMO sector and the area imaged using the AOSLO. This mask was applied to the binary CNN segmentation and the percentage of pixels containing segmented capillaries in the region covered by the mask determined the RPC density. Density was only computed if the area imaged exceeded 30% of the total area within the sector. The hemifield RPC density was calculated using a mask that covered the superior, temporal, and inferior 60° sectors to form a 180° hemifield. The mask was generated by the union of this temporal hemifield and the AOSLO imaging area. If the area covered in the imaging session was not at least 30% of the total area within the hemifield, hemifield RPC density was not computed.

### STATISTICAL ANALYSIS

For SDOCT-derived parameters, the coefficients of repeatability 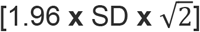 and variation [(SD/mean) **x** 100] were calculated for mean ALCSD, mean MRW, and RNFL thickness across repeated measures in each control eye. For each of these parameters, a 95% confidence interval was calculated for each control eye. For each contralateral EG eye, the first time-point of significant change in each SDOCT-derived parameter was determined as the first time-point whose value fell outside the respective 95% confidence interval and remained so at all subsequent time-points.

To determine the first significant time-point of change in RPC density, a pooled 95% confidence interval for each retinal sector was generated from the coefficient of variation obtained from repeated measures in control eyes and repeated baseline measurements in the EG eyes. As with the SDOCT-derived parameters, the first time-points of significant change in hemifield and sectoral measures of RPC perfusion parameters and RNFL thickness for each experimental glaucoma (EG) eye were determined as the first time-point whose value fell outside the pooled 95% confidence interval and had no subsequent time-points with a value that fell back within the confidence interval. RPC perfusion density and RNFL thickness were also compared over the hemifield and within each sector (superotemporally, temporally, inferotemporally).

## RESULTS

Animal characteristics and IOP data for all control and EG eyes throughout the duration of the experiment are summarized in Table 1. Mean values of IOP (± SD) across all time-points were 14.4 ± 2.2 mmHg in control eyes and 24.6 ± 4.6 mmHg in EG eyes before pharmacological intervention. The maximum IOPs measured across the control eyes and the EG eyes had mean values of 18.8 ± 2.4 mmHg and 43.4 ± 10.5 mmHg, respectively. The mean experiment duration, defined as the time between the first laser session and last AOSLO imaging session for each NHP, was 215 ± 114 days.

**Table 1.** Demographic and IOP data for all control and experimental glaucoma (EG) eyes.

| Animal ID | Age (yrs) | Gender | Mean IOP ± SD (mmHg) | | Maximum IOP (mmHg) | | Experiment Duration (Days) |
| --- | --- | --- | --- | --- | --- | --- | --- |
|  |  |  | Control Eye | EG Eye | Control Eye | EG Eye |  |
| OHT 79 | 5.5 | M | 18.2 ± 1.6 | 31.7 ± 11.1 | 22 | 54 | 350 |
| OHT 78 | 5.7 | M | 12.1 ± 2.1 | 22.2 ± 8.4 | 19 | 36 | 378 |
| OHT 80 | 5.3 | M | 13.5 ± 2.6 | 27.5 ± 14.2 | 18 | 64 | 384 |
| OHT 81 | 6.1 | F | 16.9 ± 2.2 | 23.0 ± 7.8 | 20 | 38 | 154 |
| OHT 82 | 6.1 | F | 12.9 ± 1.0 | 29.8 ± 10.9 | 15 | 43 | 182 |
| OHT 83 | 7.5 | F | 11.1 ± 1.8 | 16.0 ± 9.0 | 15 | 34 | 81 |
| OHT 84 | 7.6 | M | 15.2 ± 3.6 | 27.0 ± 17.1 | 22 | 54 | 137 |
| OHT 87 | 8.9 | F | 14.8 ± 2.7 | 23.5 ± 6.7 | 20 | 34 | 147 |
| OHT 86 | 7.9 | M | 15.5 ± 2.3 | 21.0 ± 6.2 | 18 | 34 | 119 |
| Mean ± SD | 6.6 ± 1.2 | | 14.4 ± 2.2 | 24.6 ± 4.6 | 18.8 ± 2.4 | 43.4 ± 10.5 | 215 ± 114 |

A summary of RNFL thickness (RNFLT), mean ALCSD, and mean MRW values in the control eyes of each animal over the experiment duration are presented in Supplementary Table 1. These data were used to calculate the coefficients of repeatability and variation, as previously described, for determining the first points of change in the same parameters in the fellow EG eye. The mean (± SD) values for global RNFLT, mean ALCSD, and mean MRW across all animals were 116.4 ± 6.8 µm, 200.7 ± 24.3 µm, and 340.0 ± 50.7 µm, respectively. Coefficients of repeatability and variation were small for all three of the OCT-derived parameters. The average coefficients of variation for global RNFLT, mean ALCSD, and mean MRW across animals were 5.2% ± 1.6%, 10.4% ± 3.0%, and 4.9% ± 1.2% respectively.

Longitudinal changes in SDOCT-derived parameters were observed in all EG eyes (Fig. 2). Global RNFLT and mean MRW decreased throughout the duration of the experiment in all NHPs (Fig. 2b,c). On average, across all EG eyes, the minimum values for RNFLT and mean MRW were 84.8 ± 18.2 µm and 172.3 ± 66.2 µm. These values represent a 26% and 50% decrease from baseline values of RNFLT and MRW. The anterior surface of the lamina cribrosa posteriorly deformed within each EG eye over the experiment resulting in increased values of mean ALCSD (Fig. 2d). On average, across all EG eyes, the maximum value for mean ALCSD was 420.7 ± 103.2 µm, representing a mean increase of 109% from baseline mean ALCSD levels.

**Figure 2.**
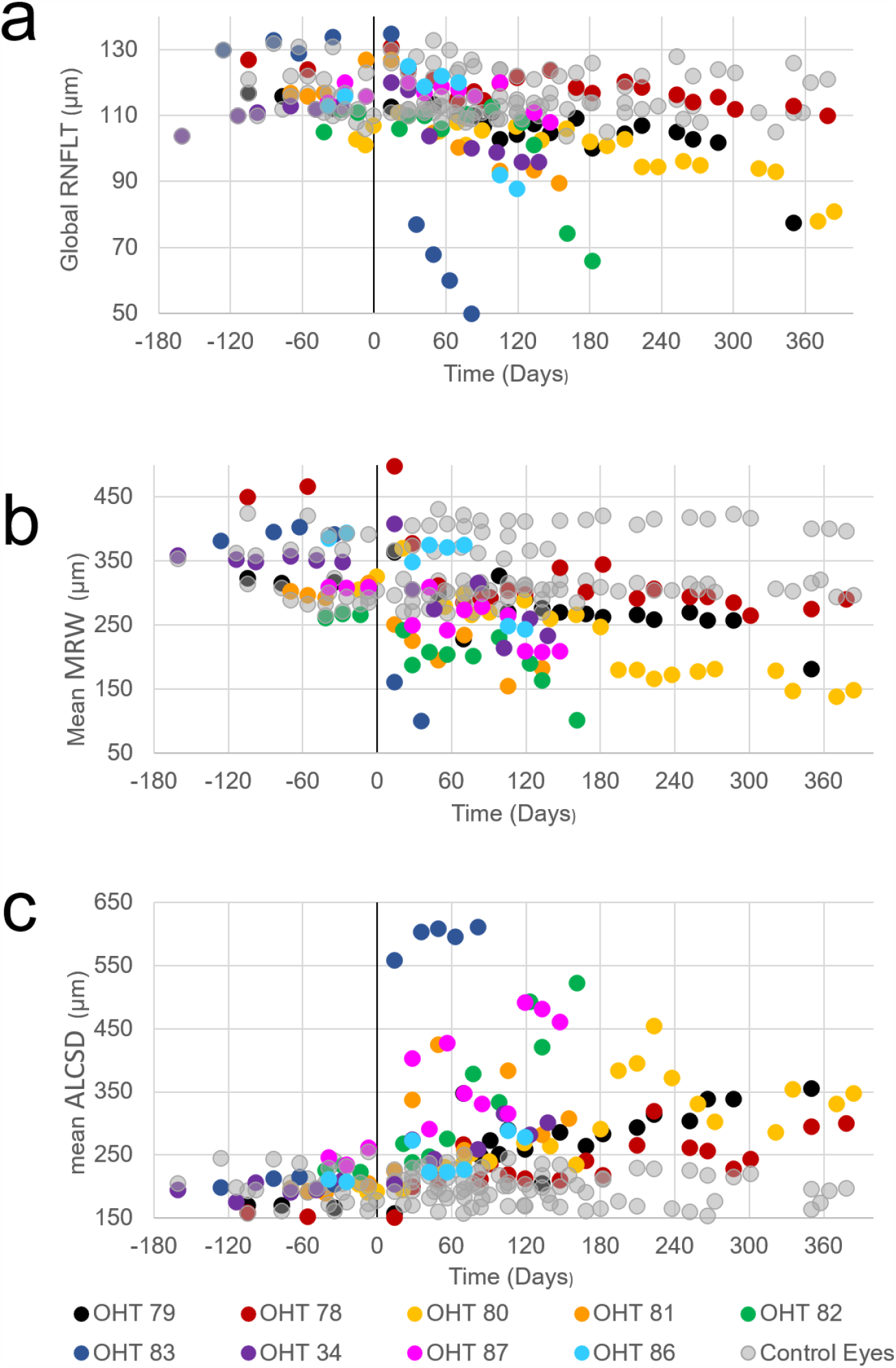
SDOCT-derived metrics of RNFLT, mean MRW, and mean ALCSD in all eyes as a function of time relative to the time of the first laser session (vertical black line at time = 0 Days) in EG eyes (filled colored circles with the legend at the bottom of the figure). Negative time-points correspond to baseline measurements (before the first laser session). Control eye data are presented as gray filled circles. After the induction of EG, (a) global RNFLT decreased, (b) Mean MRW thinned (decreased), and (c) the anterior laminar surface moved posteriorly, resulting in increased values of Mean ALCSD over the course of the experiment.

Montages of perfused retinal vasculature acquired using AOSLO imaging in a representative control eye over the duration of the experiment are shown in Figure 3. Due to experimental constraints, it was not possible to safely perform AOSLO imaging and obtain RPC data in all sectors within each control eye at all time-points. Therefore, coefficients of variation in global and sectoral values of RPC density were calculated for each animal based on repeated measures obtained over time in the animal’s control eye and repeated baseline measurements obtained in the animal’s EG eye. Mean coefficients of variation for perfused RPC density measured globally and in the superior, temporal, and inferior sectors were 19.4 ± 14.4%, 21.3 ± 15.3%, 24.3 ± 20.0%, and 31.8 ± 17.5%, respectively. Table 2 shows the baseline values of RPC density (mean ± standard deviation) and the RPC density at the time-point of first statistically significant change for each animal.

**Figure 3.**
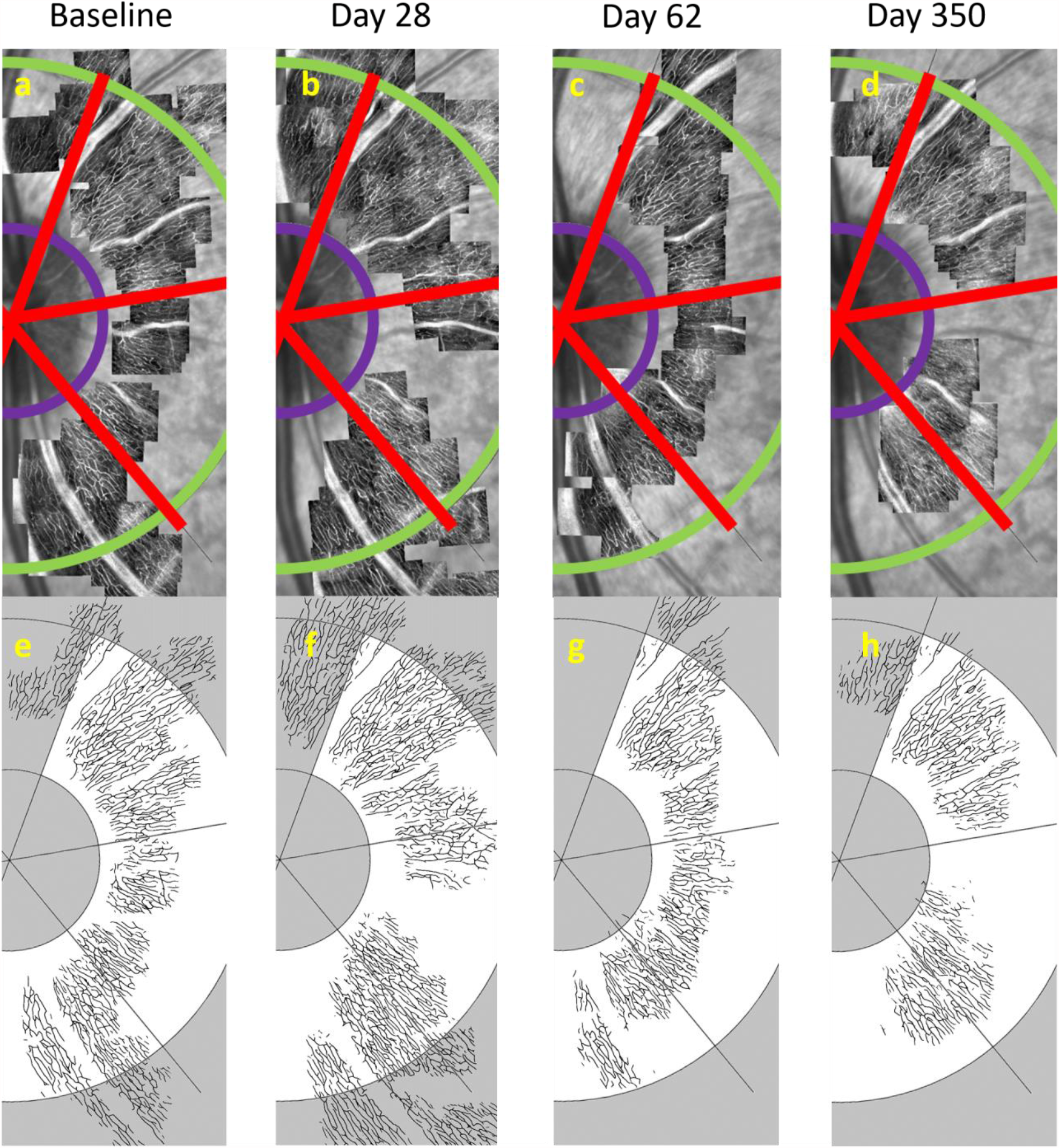
Perfused RPCs were consistently imaged in control eyes throughout the experiment. (a-d) AOSLO perfusion montages and (e-h) corresponding neural network segmentations from a representative control eye (OHT-78, OS) acquired at different time-points relative to the first laser session in the contralateral EG eye. (a) The Baseline RPC perfusion montage was overlaid on the corresponding SLO image. Red lines depict 60° sectoral boundaries for RPC analysis regions, as previously described in Figure 1. Perfused RPC density was analyzed in the annulus between the inner purple arc and outer green arc, located at radii of 3° and 8° from the center of the BMO ellipse, respectively. The corresponding CNN segmentation of the perfused RPCs illustrated in (a) is shown immediately below in (e). RPC perfusion montages acquired in the same control eye and their corresponding CNN segmentations are shown at (b,f) 28 days, (c,g) 62 days, and (d,h) nearly one year following the initial laser session performed in the EG eye.

**Table 2.**
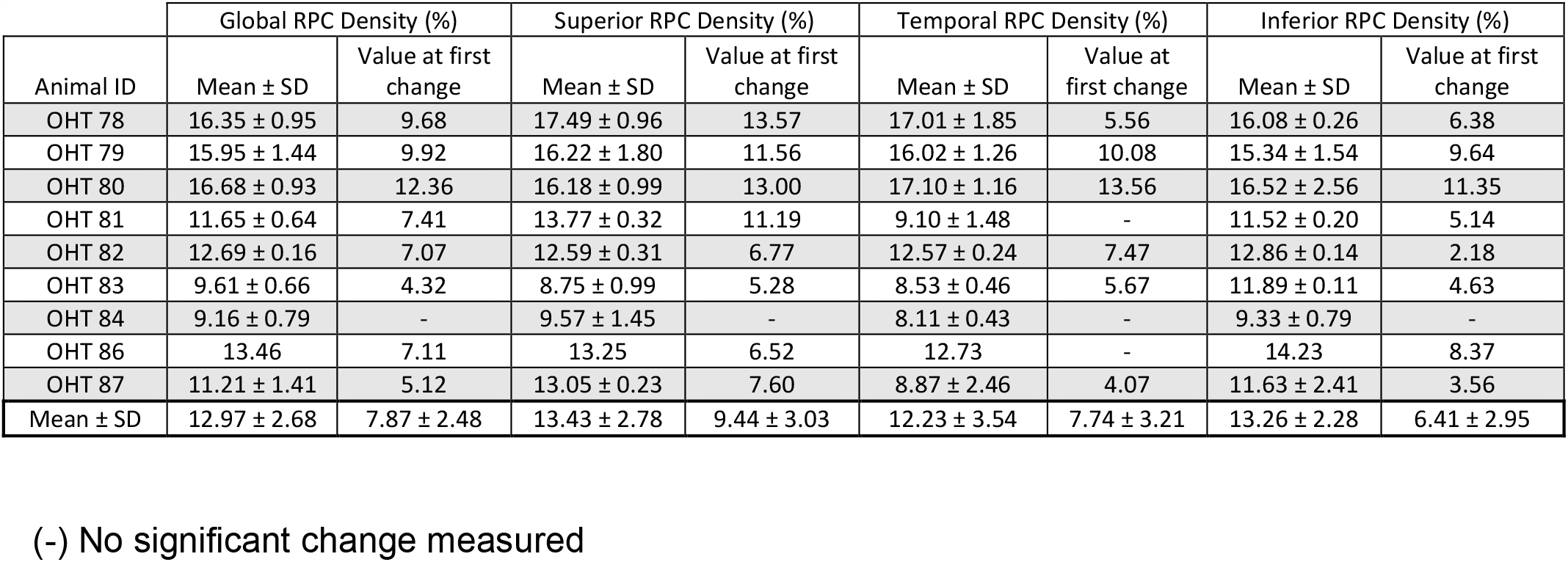
Mean values (± 1 standard deviation) of hemifield (global) and sectoral RPC densities at baseline and absolute values at the time-point of first significant change in EG eyes.

| Animal ID | Global RPC Density (%) |  | Superior RPC Density (%) |  | Temporal RPC Density (%) |  | Inferior RPC Density (%) |  |
| --- | --- | --- | --- | --- | --- | --- | --- | --- |
| | Mean ± SD | Value at first change | Mean ± SD | Value at first change | Mean ± SD | Value at first change | Mean ± SD | Value at first change |
| OHT 78 | 16.35 ± 0.95 | 9.68 | 17.49 ± 0.96 | 13.57 | 17.01 ± 1.85 | 5.56 | 16.08 ± 0.26 | 6.38 |
| OHT 79 | 15.95 ± 1.44 | 9.92 | 16.22 ± 1.80 | 11.56 | 16.02 ± 1.26 | 10.08 | 15.34 ± 1.54 | 9.64 |
| OHT 80 | 16.68 ± 0.93 | 12.36 | 16.18 ± 0.99 | 13.00 | 17.10 ± 1.16 | 13.56 | 16.52 ± 2.56 | 11.35 |
| OHT 81 | 11.65 ± 0.64 | 7.41 | 13.77 ± 0.32 | 11.19 | 9.10 ± 1.48 | - | 11.52 ± 0.20 | 5.14 |
| OHT 82 | 12.69 ± 0.16 | 7.07 | 12.59 ± 0.31 | 6.77 | 12.57 ± 0.24 | 7.47 | 12.86 ± 0.14 | 2.18 |
| OHT 83 | 9.61 ± 0.66 | 4.32 | 8.75 ± 0.99 | 5.28 | 8.53 ± 0.46 | 5.67 | 11.89 ± 0.11 | 4.63 |
| OHT 84 | 9.16 ± 0.79 | - | 9.57 ± 1.45 | - | 8.11 ± 0.43 | - | 9.33 ± 0.79 | - |
| OHT 86 | 13.46 | 7.11 | 13.25 | 6.52 | 12.73 | - | 14.23 | 8.37 |
| OHT 87 | 11.21 ± 1.41 | 5.12 | 13.05 ± 0.23 | 7.60 | 8.87 ± 2.46 | 4.07 | 11.63 ± 2.41 | 3.56 |
| Mean ± SD | 12.97 ± 2.68 | 7.87 ± 2.48 | 13.43 ± 2.78 | 9.44 ± 3.03 | 12.23 ± 3.54 | 7.74 ± 3.21 | 13.26 ± 2.28 | 6.41 ± 2.95 |

(-) No significant change measured

Longitudinal RPC perfusion images and plots of all measured parameters (Mean MRW, Mean ALCSD, global and sectoral RNFLT, global and sectoral RPC Density) are shown in Figures 4-6 for animals demonstrating different patterns of loss in RPC perfusion density and RNFLT. Figure 4 shows data from one representative animal (out of five) who demonstrated a decrease in RPC perfusion density prior to a thinning of the RNFL in its EG eye. In this animal, the initial loss of perfusion density in the inferior sector (Fig. 4b,e,j) occurred at day 14 and persisted for the remainder of the experiment. An initial decrease in RNFLT in the corresponding inferior sector was not measured until the next imaging session at day 28 (Fig. 4i).

**Figure 4.**
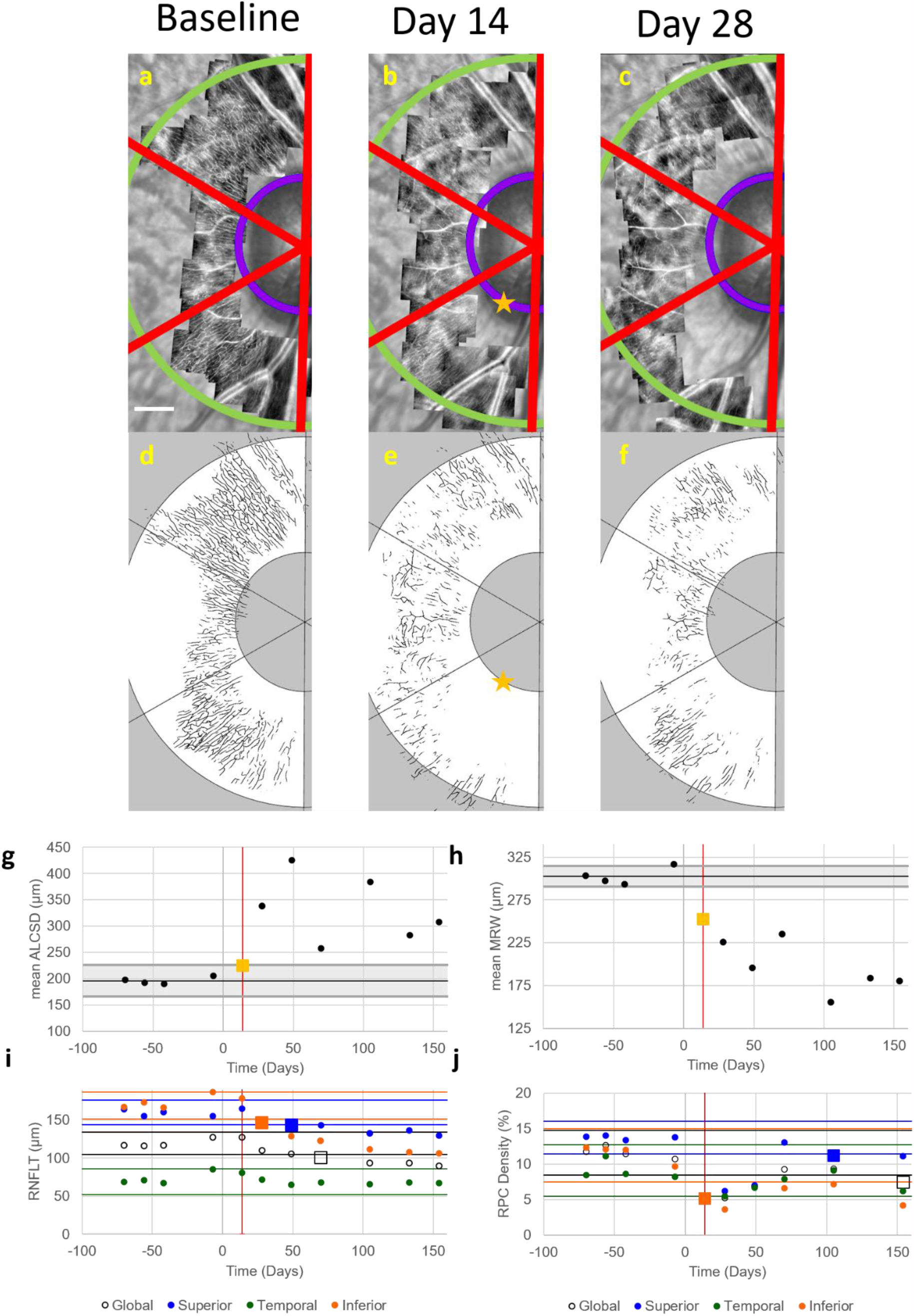
RPC perfusion density changed prior to an initial change in RNFL thickness in the EG eye of OHT-81. Images of (a-c) perfused RPCs and (d-f) their corresponding automatic segmentations acquired in the EG eye of OHT-81 at baseline (left column), the time-point corresponding to the first significant change in RPC perfusion density (middle column, 14 days after the initial laser session), and the time-point of first significant change in RNFL thickness (right column, 28 days after the initial laser session). (Note: Not all imaging time-points are included in this figure.) Perfused RPC density was analyzed globally and in 60° sectors (red boundary lines) within the annulus between 3° (inner purple arc) and 8° (outer green arc) from the center of the BMO ellipse. Scale bar in (a) represents 400 µm. The first significant change in RPC perfusion density was measured in the inferior sector at 14 days after the first laser session (yellow star). Values for (g) mean ALCSD and (h) mean MRW are plotted as a function of time for all imaging time-points before and after the time of the initial laser session (Day 0). Black horizontal lines represent the mean baseline value for these 2 parameters. Gray shaded regions represent the 95% confidence interval for each parameter calculated from data measured in the fellow control eye, with yellow squares representing the time of first significant change. Values of (i) RNFLT and (j) RPC density are plotted for global measures (black open circle) and for superior (blue circle), temporal (green circle), and inferior (orange circle) sectors at the same imaging time-points. Corresponding colored horizontal lines indicate the 95% confidence interval for each measure, with square markers representing the time of first significant change. Vertical red lines in all plots indicate the first time-point of significant change in RPC perfusion density. The first parameters to change in OHT-81 were (g) mean ALCSD, (h) mean MRW, and (j) RPC density in the inferior sector at day 14. RNFL thickness changed first in the inferior sector at day 28.

Figure 5 shows data from one representative animal (out of two) who demonstrated an initial thinning in RNFLT before an initial decrease in RPC perfusion density. Mean ALCSD was the first parameter to initially change at 49 days following the first laser session in the EG eye of OHT-79 (Fig. 5g). At 105 days after the initial laser session, there was an initial decrease in mean MRW that was accompanied by decreases in global RNFLT and the inferior sector measurement of RNFLT (Fig. 5i), as well as measurements of RPC perfusion density made globally and in the superior and temporal sectors (Fig. 5j). However, the measurements of global RNFLT, global RPC perfusion density, and superior and temporal RPC perfusion densities all returned to being within their respective confidence intervals at later time-points. The first time-point in which RPC perfusion decreased and remained outside the confidence interval occurred in the superior sector at 266 days following the first laser session (Fig. 5c,f,j).

**Figure 5.**
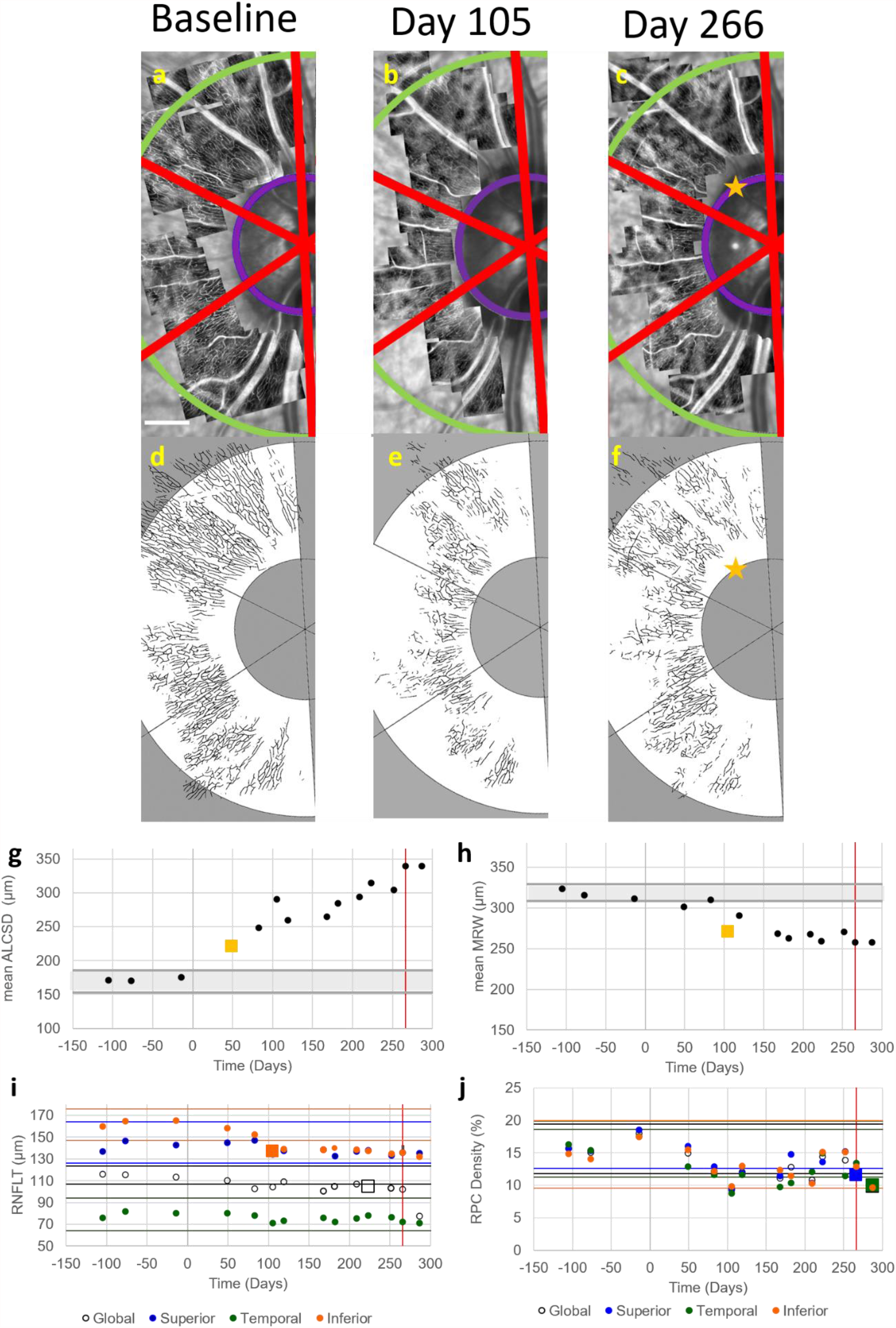
RPC perfusion density changed after the time-point of first change in RNFL thickness in the EG eye of OHT-79. Images of (a-c) perfused RPCs and (d-f) their corresponding automatic segmentations acquired in the EG eye of OHT-79 at baseline (left column), the time-point of first significant change in RNFL thickness (middle column, 168 days after the initial laser session), and the time-point of first significant change in RPC perfusion density (right column, 266 days after the initial laser session). (Note: Not all imaging time-points are included in this figure.) Perfused RPC density was analyzed globally and in 60° sectors (red boundary lines) within the annulus between 3° (inner purple arc) and 8° (outer green arc) from the center of the BMO ellipse. Scale bar in (a) represents 400 µm. The first significant change in RPC perfusion density was measured in the superior sector at 266 days after the initial laser session (yellow star). Values for (g) mean ALCSD and (h) mean MRW are plotted as a function of time for all imaging time-points before and after the time of the initial laser session (Day 0). Black horizontal lines represent the mean baseline value for these 2 parameters. Gray shaded regions represent the 95% confidence interval for each parameter calculated from data measured in the fellow control eye, with yellow squares representing the time of first significant change. Values of (i) RNFLT and (j) RPC density are plotted for global measures (black open circle) and for superior (blue circle), temporal (green circle), and inferior (orange circle) sectors at the same imaging time-points. Corresponding colored horizontal lines indicate the 95% confidence interval for each measure, with square markers representing the time of first significant change. Vertical red lines in all plots indicate the first time-point of significant change in RPC perfusion density. The first parameter to change in OHT-79 was (g) mean ALCSD (day 49), followed by (h) mean MRW (day 105). The next parameter to change was (i) RNFLT in the inferior sector (orange square, day 168), followed by the last parameter to change, (j) RPC perfusion density in the inferior sector (day 266).

Figure 6 depicts one representative case (out of two) where initial changes in RNFLT and RPC perfusion density occurred at the same imaging time-point. Nearly 100 days after the first changes were measured in mean MRW (day 21) and mean ALCSD (day 42), an initial thinning in global RNFLT and the inferior sector measurement of RNFLT occurred on day 133 (Fig. 6g). At the same time-point, a decrease in RPC perfusion density was measured globally and in all sectors (Fig. 6h).

**Figure 6.**
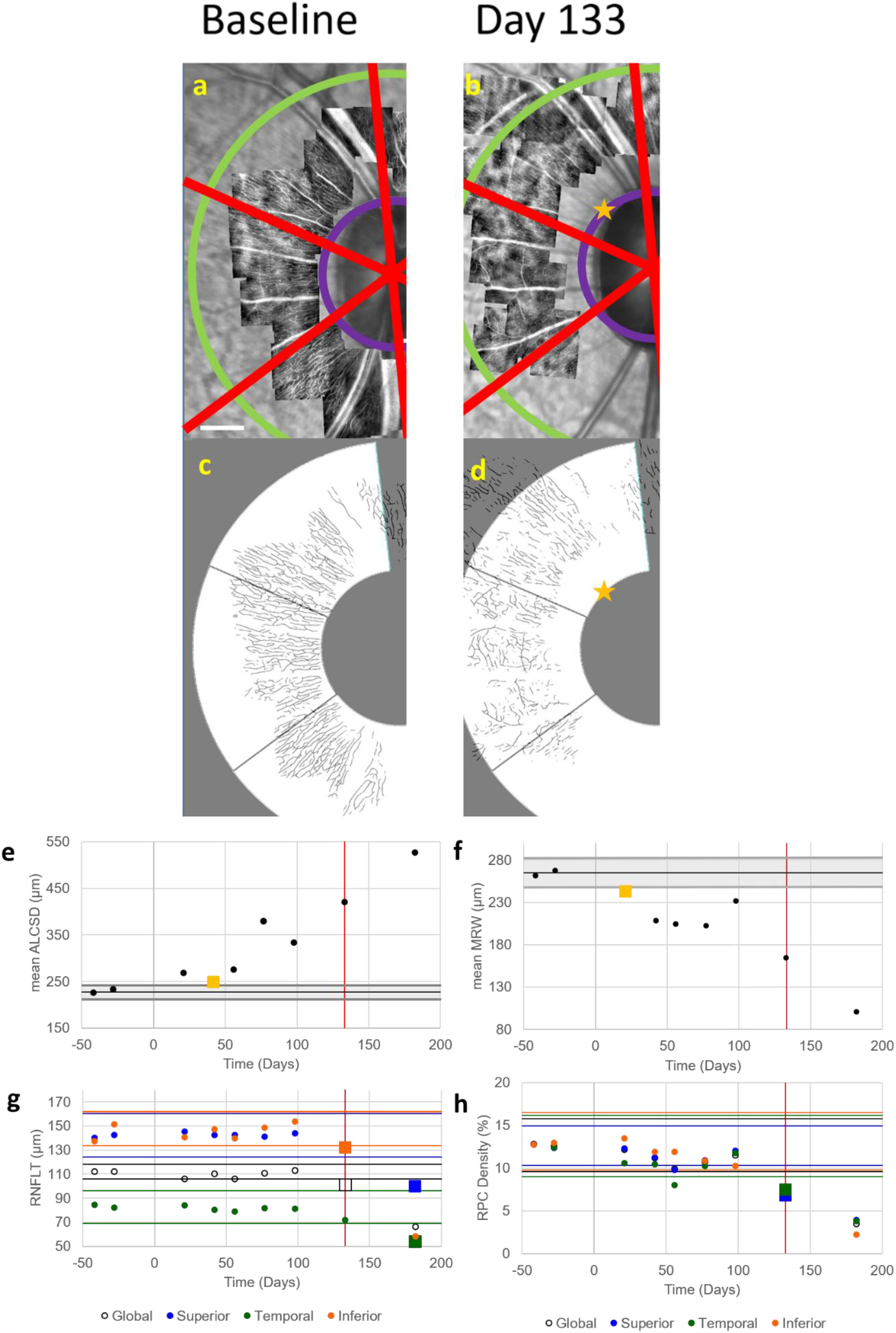
Initial changes in RPC perfusion density and RNFL thickness occurred simultaneously in the EG eye of OHT-82. Images of (a,b) perfused RPCs and (c,d) their corresponding automatic segmentations acquired in the EG eye of OHT-82 at baseline (left column) and the time-point corresponding to the first significant changes in RPC perfusion density and RNFL thickness (right column, 133 days after the initial laser session). (Note: Not all imaging time-points are included in this figure.) Perfused RPC density was analyzed globally and in 60° sectors (red boundary lines) within the annulus between 3° (inner purple arc) and 8° (outer green arc) from the center of the BMO ellipse. Scale bar in (a) represents 400 µm. The first significant change in RPC perfusion density was measured in the superior sector at 133 days after the initial-laser session (yellow star). Values for (e) mean ALCSD and (f) mean MRW are plotted as a function of time for all imaging time-points before and after the time of the initial laser session (Day 0). Black horizontal lines represent the mean baseline value for these 2 parameters. Gray shaded regions represent the 95% confidence interval for each parameter calculated from data measured in the fellow control eye, with yellow squares representing the time of first significant change. Values of (g) RNFLT and (h) RPC density are plotted for global measures (black open circle) and for superior (blue circle), temporal (green circle), and inferior (orange circle) sectors at the same imaging time-points. Corresponding colored horizontal lines indicate the 95% confidence interval for each measure, with square markers representing the time of first significant change. Vertical red lines in all plots indicate the first time-point of significant change in RPC perfusion density. The first parameter to change in OHT-82 was (f) mean MRW (day 21), followed by (e) mean ALCSD (day 42). Initial decreases in global and sectoral values of RPC perfusion density and RNFLT occurred simultaneously at day 133.

The first time-points of change in all ONH and RPC parameters on global or sectoral levels are illustrated in Figure 7 for each EG eye as a function of study time relative to the date of the first laser session. An increase in mean ALCSD was among the first structural changes in 7 of 9 eyes. A simultaneous decrease in mean MRW occurred in 6 of these 7 eyes, and 2 also exhibited a decrease in RPC perfusion density. Mean MRW was the first parameter to solely change in one EG eye (OHT 82) while RPC density solely changed first in the 9th EG eye (OHT 86). All EG eyes experienced decreases in RPC perfusion density and RNFLT throughout the duration of the experiment. A decrease in RPC perfusion density first occurred prior to the first instance of RNFL thinning in 5 of 9 EG eyes (OHT-78, 80, 81, 83, 86). The first decrease in RPC perfusion density and RNFLT occurred simultaneously in 2 EG eyes (OHT-82, 87), while an initial thinning of the RNFL occurred before an initial decrease in RPC perfusion density in 2 EG eyes (OHT-79, 84).

**Figure 7.**
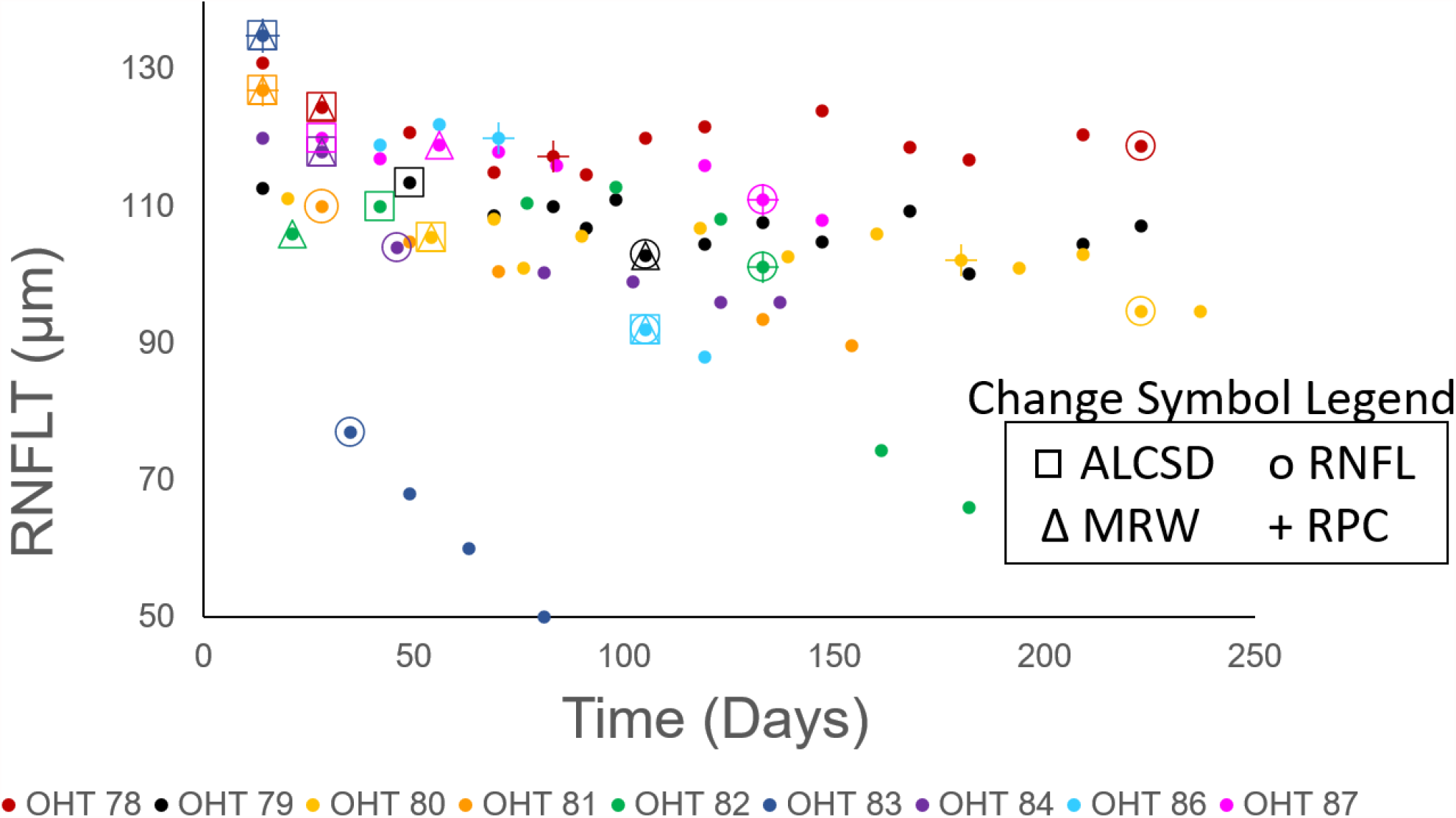
Time of first significant change measured in all parameters for each EG eye. RNFLT values are plotted as a function of study time for all EG eyes (filled circles, with each color representing a different EG eye). The time-points corresponding to the first significant change in Mean ALCSD, Mean MRW, RNFLT, and RPC density on a global or sectoral level are shown using a square, triangle, circle, and cross, respectively. A progressive decrease in global RNFLT was measured throughout the course of the study for all EG eyes. An increase in mean ALCSD (square) was the first structural change to occur in 7 of 9 EG eyes. A change in mean MRW (triangle) and RPC perfusion density (“+”) simultaneously accompanied this first change in 6 EG eyes and 2 EG eyes, respectively. A decrease in RPC perfusion density was the first structural change to solely occur in one EG eye (OHT 86). The first change in RPC perfusion density occurred prior to the first change in RNFLT (circle) in 5 of 9 EG eyes.

## DISCUSSION

The purposes of this study were to (1) characterize the time-course of *in vivo* changes in RPC perfusion concurrent with changes in ONH structure and RNFL thickness and (2) to determine whether changes in RPC perfusion occur prior to changes in RNFL thickness in early experimental glaucoma. ONH parameters (mean ALCSD, mean MRW) changed before retinal parameters (RPC density, RNFLT) in all eyes but one, in which RPC density was the first parameter to change. RPC density significantly decreased before RNFL thinning occurred in the majority of eyes studied.

The *in vivo* IOP and ONH measurements found in healthy control and EG eyes in this study are similar to values published from other studies. The average maximum IOP measured across all EG eyes in this study was 43.4 ± 10.5 mmHg, which was 231% greater than the average maximum IOP across the contralateral control eyes of 18.8 ± 2.4 mmHg. Work from previous studies using the same or a similar method of inducing glaucoma experimentally in NHPs have resulted in a +282%,^35^ +280%,^46^ and +241%^47^ difference in the mean maximum IOP between EG and control eyes. For ONH measures, the average mean ALCSD for all control eyes across all time-points in this study was 200.7 ± 24.3 µm, which is similar to values of mean ALSCDs reported previously in healthy NHP eyes (ranging from 214 – 230 µm).^35,48–50^ The average mean MRW for all control eyes across all time-points in this study (340.0 ± 50.7 µm) is also similar to the mean value previously reported in control eyes by Ivers et al.^35^ (308.7 ± 55.1 µm) and is within the range reported by Strouthidis et al. (∼200 - 425 µm).^44^ In addition, the mean global RNFLT measured in the control eyes over the course of this study (116.4 ± 6.8 µm) is within the range of values of global RNFLT published in healthy NHP eyes (∼101 – 124 µm).^35,39,51,52^

Coefficients of variation (CoVs) computed using the control eye data for all OCT-derived parameters in this study were also consistent with CoVs for the same metrics reported by other studies. The CoVs for mean ALCSD, mean MRW, and global RNFLT in this study were 10.4 ± 3.0%, 4.9 ± 1.2%, and 5.2 ± 1.6%, which are consistent with previously reported CoVs for mean ALCSD (4.5%),^35^ mean MRW (2.3% to 11.9%),^35,39^ and global RNFLT (1.7% to 7.5%).^35,39^ The low CoVs found in this study for the OCT-derived metrics and their similarity to values acquired in other studies lend support to the consistency of the experimental methods implemented in this study.

This study reports the first values of RPC perfusion density near the ONH in healthy and diseased NHP eyes based on images obtained using split detector AOSLO imaging to our knowledge. The values for RPC perfusion density measured from AOSLO images acquired in healthy NHP retinas in this study (13.06 ± 2.87 %) are lower than those that have been reported when using OCTA imaging in the NHP eye (35.4 ± 3.4%).^53^ A combination of factors could potentially contribute to differences in RPC perfusion density values between these two imaging modalities. For example, AOSLO images have better resolution than OCTA images (due to their correction of the eye’s higher order aberrations), and have been shown to closely agree with histological images of retinal vasculature.^54^ The decreased lateral resolution inherent in OCTA perfusion images yields a broader point (or line) spread function and results in larger diameter vessels, which can lead to the calculation of higher perfusion densities. In addition, a typical OCTA image represents a maximum intensity projection from an OCTA volume scan. Therefore, OCTA images usually include vasculature being perfused at multiple depths within the slab being quantified. Conversely, AOSLO imaging typically collects light over a smaller depth. Consequently, it is likely that less perfused vasculature will be observed in an AOSLO perfusion image over a comparably sized area (relative to an OCTA perfusion image) and yield decreased perfusion density values.

To our knowledge, we also report the first CoV values for quantifying RPC perfusion density near the ONH in healthy eyes in images obtained using split detector AOSLO imaging. The CoVs for RPC perfusion density in each sector ranged from 21.0 – 26.5%. Several factors could contribute to the variability of the perfusion density measurements. For each AOSLO imaging session, the plane of best focus for RPCs is manually selected after using the confocal AOSLO channel to section through the retina and visualize the most superficial retinal nerve fiber bundles. Since the retina is a curved surface and retinal thickness is uneven around the ONH, the focus must be checked throughout the imaging session, as the curvature and thickness of the retina can change dramatically near the rim of the ONH. An additional factor that could impact the CoV is the size of the field of view of the image. Lee et al. evaluated the CoV for macular vessel density using OCTA images of different field sizes and found that the CoV for vessel density was much greater when using a small, 1mm x 1mm scan (18.552%) compared to a larger, 6mm x 6mm scan (4.042%).^55^ The smaller field size of the individually acquired AOSLO images (∼0.6mm x 0.6mm field of view) could contribute to a larger CoV. In addition, differences in the areas that were imaged between sessions could impact the CoVs calculated in this study. Despite our best attempts to image exactly the same areas in each eye across all time-points, slight differences occurred in the portions of the retina that were imaged within each eye over time. Most AOSLO systems do not currently possess an inbuilt eye tracking system, as is common on most OCT systems, making it more challenging to consistently position the animal’s head and examine the same retinal locations between sessions. Also, the AOSLO system used in this study relies on post-processing to stabilize AOSLO videos and register and align its images relative to a reference frame. Previous work has shown that the selection of a reference frame with minimal distortion is important to achieving high-quality stabilization and optimal image quality in adaptive optics^56^ and OCT imaging.^57^ Automating the focus selection and adding eye tracking capabilities to the AOSLO system could potentially decrease variability in the areas imaged between sessions and result in smaller confidence intervals for RPC perfusion density.

Mean ALCSD and/or mean MRW were the first measured parameter to change in 8 of 9 animals, while no NHPs had significant RNFL thinning prior to a change in any ONH parameter. These findings are consistent with other studies that have examined ONH parameters and RNFLT and found that ONH-related parameters tend to change before retinal parameters.^35,48^ An initial change in RPC perfusion density occurred simultaneous with an initial change in an ONH parameter in 2 of the 8 aforementioned eyes, while also occurring solely on its own in the 9^th^ eye (i.e., in OHT 86, prior to a measured change in any ONH parameter in that eye). The first change in RPC perfusion density was measured simultaneous to or following a first change in an ONH parameter in 8 of 9 EG eyes. Future experiments could examine the extent to which a loss in RPC perfusion is a more primary mechanism of glaucoma in some eyes, or possibly occurs secondarily as a consequence of ONH changes in other eyes.

Decreases in RPC perfusion could potentially decrease the supply of nutrients to and the removal of waste from RNFL axons in the region of perfusion loss. Therefore, a local loss in RPC perfusion could be expected to result in a subsequent loss in RNFL axons (or a decrease in RNFLT) in the corresponding local area. Such local relationships were observed in some EG eyes in this study. For example, an initial loss in RPC perfusion density in the inferior sector was observed within the EG eye of OHT 81 (Fig. 4j, day 14) before an initial thinning of the RNFL in the corresponding inferior sector (Fig. 4i, day 28). Similarly, RPC perfusion density decreased in the superior sector within the EG eye of OHT 86 at 70 days after the initial laser session, which was prior to the first significant decrease measured in RNFLT in the corresponding superior sector at 105 days following the initial laser session (data not shown). In these examples, the time course of alterations in RPC perfusion density and RNFLT may indicate that a loss of perfusion resulted in subsequent damage to RNFL axons in corresponding locations and an ultimate decrease in RNFLT. However, mixed results were found in other EG eyes. For example, while initial decreases in RPC perfusion density were measured in the superior and temporal sectors of OHT 82 (Fig. 6h, day 133) before an initial thinning of the RNFL in the corresponding superior and temporal sectors (Fig. 6g, day 182), an opposite relationship was found in the inferior sector of the same eye, in which RNFLT decreased (day 133) before an initial decrease in RPC perfusion density (day 182). In another example, RPC perfusion density first decreased in the temporal sector within the EG eye of OHT 78 at 83 days after the initial laser session (data not shown) while a change in RNFLT relative to baseline was never measured in the corresponding temporal sector.

Different reasons could account for the different patterns of loss in RPC perfusion density and RNFLT in the EG eyes examined in this study. First, it is quite possible that RNFL axons have different susceptibilities to changes in vascular perfusion in different eyes. In addition, despite our best efforts to consistently image the same retinal regions, it also possible that alterations in RPC perfusion occurred outside of the regions that were imaged at a given time-point or outside of the annulus analyzed in this work. Furthermore, these observed patterns were influenced by our method for determining the time-point of first change (i.e., the time when a given parameter first fell outside of its confidence interval limits *and* remained outside of these limits). Different results could have been found if we only considered the time-point at which a given parameter first fell outside of the confidence interval and did not require that the parameter remain outside of the interval. For example, RPC perfusion density measured in the superior sector within OHT 81 (Fig 4j) first fell outside of its confidence interval at day 14, but rebounded back within the confidence interval at a later time-point (day 70) before dropping back outside and remaining outside of the confidence interval at day 105. Due to our more conservative criterion (i.e., requiring the parameter to also remain outside of the confidence interval), the reported time of the first change was 105 days. However, without this criterion, the initial change would have occurred at day 14, which would have been prior to the first measured change in RNFLT in the superior sector (Fig. 4i) at day 49. One potential reason as to why RPC perfusion density would decrease and later increase could be that the capillaries imaged at later time-points contained both RPCs and deeper capillaries that were subsequently visible due to losses in overlying RPCs. While great effort was taken to ensure that only the most superficial capillaries were imaged at each time-point, it is possible that the most superficial capillary network that was visible at later time-points in an EG eye with a reduced RNFL could have been a deeper capillary plexus. In addition, it is important to recall that perfusion images only show vessels that are actively perfused. If capillaries are non-perfused due to an ischemic event or blockage upstream of the capillary blood supply that later resolves, capillaries could reperfuse and be imaged at a later time.^58^ Decreased autoregulation of the capillary network could also result in transient perfusion decreases. It has been shown (using Doppler laser flowmetry) that there is decreased autoregulation of blood flow in the major vasculature in the ONH in eyes with glaucoma.^59^ Some eyes could be more susceptible to decreased vascular autoregulation and vascular instabilities could be more prominent in these eyes, leading to alterations in perfusion over time.

In summary, we longitudinally examined the time-course of earliest change in radial peripapillary capillaries in a monkey model of experimental glaucoma. The results from this study show that RPCs change prior to a decrease in RNFLT in the majority of eyes with experimental glaucoma. These data suggest that vascular factors may make some eyes more susceptible to axonal loss in early stages of the disease. Future work could examine changes in RPC perfusion and structure in tandem with assessments of blood pressure, intracranial pressure and ocular perfusion pressure to better understand vascular dynamics and their implications in the development and progression of glaucoma.

## ACKNOWLEDGEMENTS

Research was supported by BrightFocus Foundation National Glaucoma Research Grant (G2018061), NIH Grants R01 EY029229 and P30 EY007551, and the University of Houston College of Optometry. The authors thank Ron Harwerth for helpful discussions and guidance during the NHP experiments. The authors acknowledge the use of the Maxwell/Opuntia/Sabine Clusters and the advanced support from the Research Computing Data Core at the University of Houston.

---

**Supplementary Table 1.**
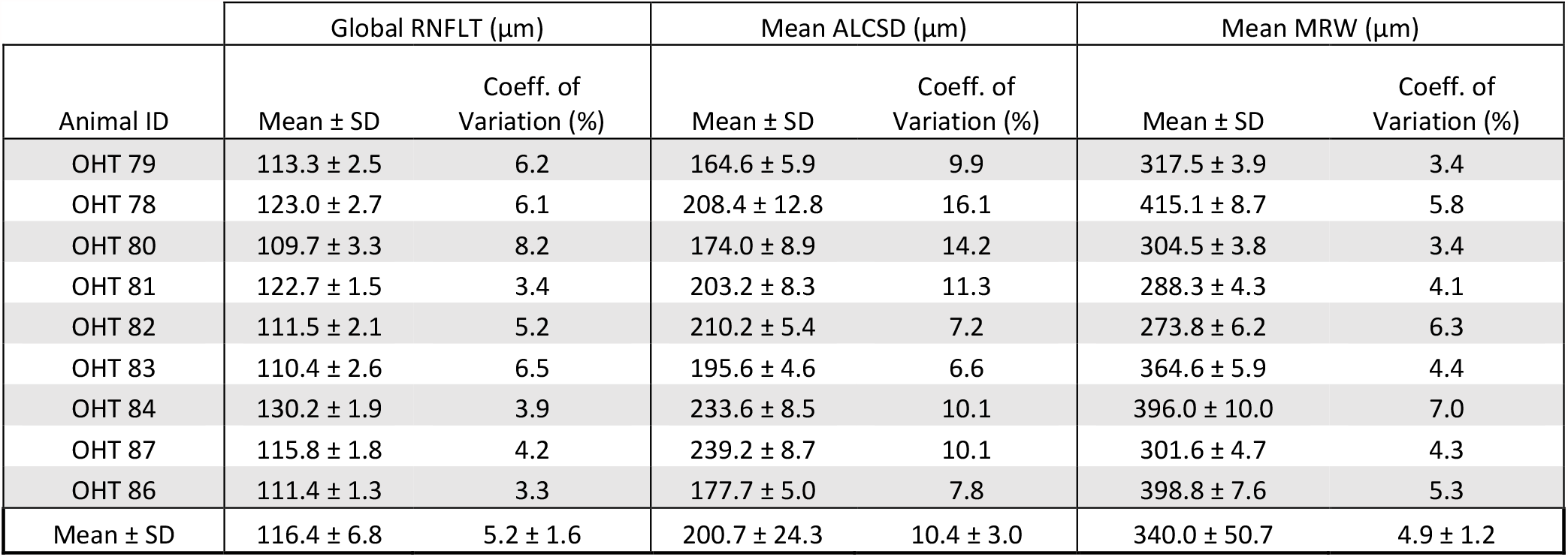
OCT-derived RNFLT, mean ALCSD, and mean MRW values across all study time-points for each control eye.

